# Development and validation of methods to assess red crown rot (*Calonectria ilicicola*) severity in soybean: standard area diagram set for roots and diagrammatic scale for canopy

**DOI:** 10.64898/2026.08.11.744294

**Authors:** Boris X. Camiletti, Juan A. Paredes, Bruno D. Pugliese, N. Dennis Bowman, Darcy E. P. Telenko, Carl A. Bradley

## Abstract

Red crown rot of soybean (RCR), caused by *Calonectria ilicicola*, is an emerging soilborne disease whose quantification is challenging due to its complex symptom development across root and foliage levels. This study developed and evaluated a multi-scale framework to improve the assessment of RCR severity from controlled environments to field conditions using root imaging and standardized visual scales. Under controlled conditions, a standard area diagram (SAD) for root necrosis was developed and validated, and SAD-assisted evaluations significantly improved accuracy, precision, and inter-rater agreement compared with unaided assessments. In field conditions, a diagrammatic symptom scale (DSS) was developed using consensus-rated images from experts and showed high reliability, repeatability, and reproducibility across 18 raters, with strong intra- and inter-rater agreement. This study developed and evaluated complementary methods to improve the assessment of RCR severity from controlled environments to field conditions using root imaging and standardized visual scales.

## 1. Introduction

Red crown rot (RCR) of soybean (*Glycine max*) is a recently emerging soilborne disease in the U.S. Midwest caused by *Calonectria ilicicola*, a pathogen that infects plants primarily through the root system (Bonkowski et al. 2024; Kleczewski et al. 2019; Neves et al. 2023). Infection often begins early in the growing season, but symptom expression can be delayed, complicating disease detection and severity assessment (Jiang and Xie 2023; Kim et al. 1998; Kuruppu and Russin 2004). Characteristic belowground symptoms include progressive root necrosis and loss of lateral roots, which can result in reduced plant size and, in extremely severe cases, stand reduction shortly after planting. As infection progresses during vegetative stages, reddish to dark brown discoloration may develop at the lower stem near the soil line, where reproductive structures (perithecia) can form during reproductive stages under favorable conditions (Kleczewski et al. 2023). Typical canopy expression includes interveinal chlorosis, necrosis, wilting, and general plant decline (Kleczewski et al. 2023; Roy 1989). Foliar symptoms most commonly develop during reproductive stage, although their onset and intensity can vary considerably among environments (Kleczewski et al. 2023). Because substantial root damage can occur before canopy symptoms become visible, plants may experience physiological stress and yield penalties even in the absence of clear foliar expression (Paredes et al. 2026), complicating disease evaluation in field trials.

Short-term experiments conducted under greenhouse or growth-chamber conditions provide a rapid and controlled framework to evaluate soybean responses to *C. ilicicola* infections, particularly during early seedling stages when root colonization and necrosis are most pronounced (Paredes et al. 2026). These short-term assays are especially well suited for differentiating seed treatment performance and have become a practical approach for screening fungicide and biological seed treatments intended to protect the root system against the pathogen. To support disease quantification in such controlled assays, Jiang et al. (2020) proposed a qualitative ordinal scale ranging from 0 to 5, based on visual assessment of root necrosis and seedling viability, where 0 represents no visible symptoms, scores 1 to 4 represent progressively increasing disease severity, and 5 corresponds to complete seedling death. This scale has been widely adopted in resistance screening studies because it is rapid, intuitive, and captures biologically meaningful stages of disease progression during early plant development (Antwi-Boasiako et al. 2024). However, as an ordinal scale, it summarizes symptom severity into discrete categories, which may limit separation when differences among treatments are subtle. Quantitative approaches for disease severity assessment may help to overcome this limitation.

Under field conditions, RCR produces foliar symptoms that closely resemble those of sudden death syndrome (SDS; caused primarily by *Fusarium virguliforme*), particularly the interveinal chlorosis and necrosis. Since SDS has been extensively studied and standardized field rating procedures are well established (Brown et al. 2023; Gongora-Canul et al. 2012; Kandel et al. 2020; Raza et al. 2020), its foliar assessment framework has frequently served as a reference for evaluating similar symptom expression. The SDS scoring system integrates disease incidence and severity into a disease index and employs a 0 to 9 ordinal scale that reflects increasing chlorosis, necrosis, progressive defoliation, and ultimately premature plant death. Despite these visual similarities, important biological differences exist between the two diseases. RCR typically induces wilting and canopy decline but does not cause the pronounced premature defoliation characteristic of SDS (Kleczewski et al. 2023). Because defoliation defines the upper categories of the SDS severity scale, direct application of this system may not accurately represent RCR symptom progression and could lead to misclassification of advanced stages. Therefore, adaptation of the SDS-based framework is necessary to accurately capture RCR foliar expression while maintaining a structured and reproducible assessment approach.

RCR is expressed at multiple biological scales throughout disease development, making no single assessment method suitable for all experimental contexts. During early infection under controlled conditions, root severity provides the most direct measure for disease severity and treatment efficacy assays. In contrast, disease impact in the field is reflected by progressive canopy symptoms that require standardized visual assessment. Therefore, our objectives were to i) develop and validate a root SAD for quantitative assessment of RCR severity under greenhouse conditions; (ii) develop and standardize an adapted severity scale for canopy field evaluations of RCR. By integrating controlled-environment assessment tools and a standardized field foliar severity scale, this study aimed to establish a comprehensive multi-scale framework for quantifying RCR severity and progression under both experimental and field conditions.

## 2. Materials and Methods

### 2.1. SAD for root rot severity

#### 2.1.1. SAD development

A total of 66 individual 12 day-old root images, representing a wide range of levels of root rot severity, were collected from soybean plants grown under a gradient of *C. ilicicola* inoculum (0.25%, 0.5%, 1% v/v) in a growth chamber experiment described by Paredes et al. 2026. Briefly, soybean cultivar Xitavo XO3922E (BASF, Research Triangle Park, NC, USA) was grown in pots filled with general-purpose potting mix soil (1:1:1 (v/v/v) mixture of soil, peat, and perlite). *C. ilicicola* inoculum was produced on colonized sorghum grain following Kleczewski et al. (2023) using two isolates from Illinois: NRRL 64312 (CC1) and 22-4A3 (MAD). The inoculum was incorporated into the soil at 0.25%, 0.5%, and 1% (v/v) before planting. Plants were maintained under a 12-hour light/12-hour dark cycle (dual-lamp fluorescent lighting, approximately 3,678 lux) at 24±1°C in a growth chamber (Thermo Scientific Precision, PR505755L). Soybean plants were harvested 12 days after emergence, and roots were gently washed under running water to remove all soil residues. Cleaned roots were placed on absorbent paper towels, then positioned in a standardized photo studio light box (60 × 60 × 60 cm) and photographed using a smartphone camera under consistent conditions, with roots placed against a white paper to maximize contrast between roots and the background.

Each image was stored in JPG format, and the percentage of root rot affected were quantified using ‘Pliman’ (Olivoto, 2022) R package, which automates image analysis by applying segmentation methods to differentiate between diseased and healthy tissue. Image scale was standardized using the ‘dpi()’ function prior to image analysis, based on a reference square with known dimensions included during image acquisition for scale calibration. Individual roots were segmented from the background using the ‘analyze_objects()’, and the total root area was extracted using ‘get_measures()’ function, with the appropriate DPI specified for scale calibration. Diseased areas were obtained using the ‘measure_disease()’ function, calibrating color thresholds for healthy tissue, symptomatic tissue, and background. The segmentation accuracy was visually confirmed, and the percentage of diseased tissue relative to the total root area was calculated to obtain the ‘actual severity’ for each root image.

The root SAD was developed using six representative images corresponding to increasing levels of root necrosis, selected to depict the range of disease severity observed in the study.

#### 2.1.2. SAD validation

A set of 60 roots images were available for the validation process, after excluding the six representative roots images used to construct the SAD. The images were displayed on a computer screen. A total of 15 raters participated in the SAD validation. All raters were researchers with backgrounds in disease assessment, and each one evaluated the complete set of images in sequence. The evaluation process began with an initial round of severity estimation without SAD assistance, followed by a round using the SAD provided for the entire images set. Raters were instructed to estimate the percentage of diseased root area for each image. When using the SAD, they were directed to use it as a visual reference, comparing each root image to the diagram that most closely matched its affected area, using this comparison to guide their percentage-based severity estimates (James 1971; Del Ponte et al. 2017).

For each rater, the mean and absolute error was calculated as the difference between the estimated and actual severity. Lin’s concordance correlation coefficient (Lin’s CCC), the generalized bias coefficient (C*b*), and precision (Pearson’s *r*) were calculated using ‘epi.ccc’ function from the ‘epiR’ R package (Madden et al. 2017; Del Ponte et al. 2017)

To quantify the effect of the SADs, paired comparisons between unaided and aided assessments were performed for each evaluation metric across raters. For each metric, paired differences (aided - unaided) were calculated at the rater level, and 95% confidence intervals for the mean differences were estimated using 2,000 bootstrap resampling. Statistical inference was based on whether the confidence intervals included zero (α = 0.05) (Brás et al. 2020; Moreira et al. 2019; Del Ponte et al. 2017).

#### 2.1.3. Comparing the use of SAD to ordinal qualitative scale

The same set of root images was classified by the raters using the scale proposed by Jiang (2020), (hereafter referred to as “ordinal scale”) an ordinal-qualitative scale ranging from 0 to 5. The six-point rating scale was as follows: 0 = no visible symptoms, 1 = small brown necrotic lesions on the primary root, 2 = brown necrotic lesions extending all over the primary root and some lateral roots, 3 = root rot became evident with 50% root-loss by rot and severe brown necrosis in the underground stem, 4 = almost all roots rotted and lost, 5 = seedlings are dead.

To compare consistency among assessment methods (unaided, SAD-assisted, and Jiang’s score), inter-rater agreement was evaluated using the intraclass correlation coefficient (ICC) using the ‘icc’ function from the ‘irr’ R package (Gamer et al., 2019). Overall Concordance Correlation Coefficient (OCCC) was calculated for the unaided and SAD-assisted assessments using the function ‘epi.occc’ from epiR package (Stevenson and Sergeant, 2025) for evaluating agreement among multiple raters (Del Ponte et al. 2017).

Additionally, a subset of root images was selected based on observed disease severity and assigned to three synthetic treatment groups (20 root images per treatment). These simulated treatments corresponded to necrosis mean values of A = 8.3, B = 41.2, and C = 63.3, which were significantly different according to Tukey’s test (α = 0.05). To evaluate the ability of each assessment method to discriminate among severity levels, one-way ANOVA was performed separately for each rater using SAD-assisted, unaided, and proportional odds logistic regression was used for the Jiang’s score-based evaluations.

### 2.2. Development of a diagrammatic symptom scale (DSS) for canopy disease severity

#### 2.2.1 DSS development

Soybean plants at R4–R5 growth stages exhibiting RCR symptoms were selected for image acquisition from fields with active RCR epidemics during the 2025 growing season. To ensure that disease assessments reflected typical field scouting conditions, symptomatic plants were visually isolated at the moment of photography while remaining in their original position within the crop canopy and surrounded by neighboring plants. This approach preserved the natural field context in which disease severity is commonly evaluated.

A total of 74 images covering a wide range of symptoms were collected. From this pool, one representative image for each disease severity score (0–9) was selected based on expert interpretation and used to construct the RCR diagrammatic symptom scale (RCR-DSS). The scoring framework was adapted from the established SDS foliar severity scale, modifying it to better reflect RCR symptom expression while preserving the 0–9 ordinal structure, with representative images serving as visual references to guide severity estimation across different levels of symptom expression.

All the sets of images were independently scored by four experts (with extensive experience in soybean disease and familiar with RCR), using the developed RCR-DSS. A consensus severity score was assigned to each image and used as a reference when at least three of the four experts agreed on the same severity class. This consensus score was used as the “true score” for subsequent validation analyses. Images reaching consensus were compiled into a final dataset, resulting in 53 images that were assembled into a PDF file for further validation analyses.

#### 2.2.2. DSS validation

The RCR-DSS was validated using a multi-rater, real image-based assessment framework. Because the scale is an ordinal descriptive scale, no unaided control was included. Therefore, the evaluation does not quantify improvement attributable to the DSS itself, but rather assesses the reliability, agreement, and reproducibility of severity estimates obtained using the proposed RCR-DSS.

A total of 18 raters with backgrounds in disease assessment independently evaluated the 53-image set representing the full range of RCR severity under natural infection conditions. Before scoring, raters were instructed to read the description associated with each severity category. Each rater then evaluated the complete image set displayed on a computer screen and assigned a severity score from 0 to 9 using the RCR-DSS as a visual reference. Each rater completed two independent evaluation rounds in separate sessions to assess scoring consistency.

Agreement and accuracy were calculated by comparing each rater’s scores against each consensus reference value, and quantified using root mean squared error (RMSE, overall deviation from consensus), mean error (systematic over- or underestimation), and Lin’s concordance correlation coefficient (LCCC) calculated using ‘epi.ccc’ function from the ‘epiR’ package. Additionally, individual deviations were calculated as the difference between each rater’s score and the consensus (group mean) score per image.

Intra-rater repeatability was assessed by comparing scores across the two rounds. Agreement was quantified using quadratic weighted Cohen’s kappa. This metric reflects the consistency of the raters when scoring the same image at different times. Differences between rounds were evaluated using paired comparisons for RMSE and mean error, with 95% confidence intervals obtained by 2,000 bootstrap resamples. Results were considered significant when confidence intervals did not include zero (α = 0.05).

Inter-rater reproducibility was assessed using the intraclass correlation coefficient (ICC) from a two-way random-effects model with absolute agreement. ICCs were calculated for each round and for the pooled dataset.

## 3. Results

### 3.1. SAD for root rot severity

#### 3.1.1. SAD development

The 66-roots image set covered the full severity range and included 18 roots in the 0– 20% range, 12 in the 20–40% range, 16 in the 40–60% range, 11 in the 60–80% range, and 5 in the 80–100% range. The SAD was designed considering the observed minimum and maximum severities, with severity levels distributed across the observed range following best practices for diagrammatic scale construction (Del Ponte et al., 2017). The root SAD consisted of six true-color images representing severity levels of 0, 10, 30, 50, 70, and 90% root rot (Figure 1). These images provided a semi-continuous linear percentage scale spanning the observed severity range, from healthy roots (0%) to severe disease (90%).

**Figure 1.**
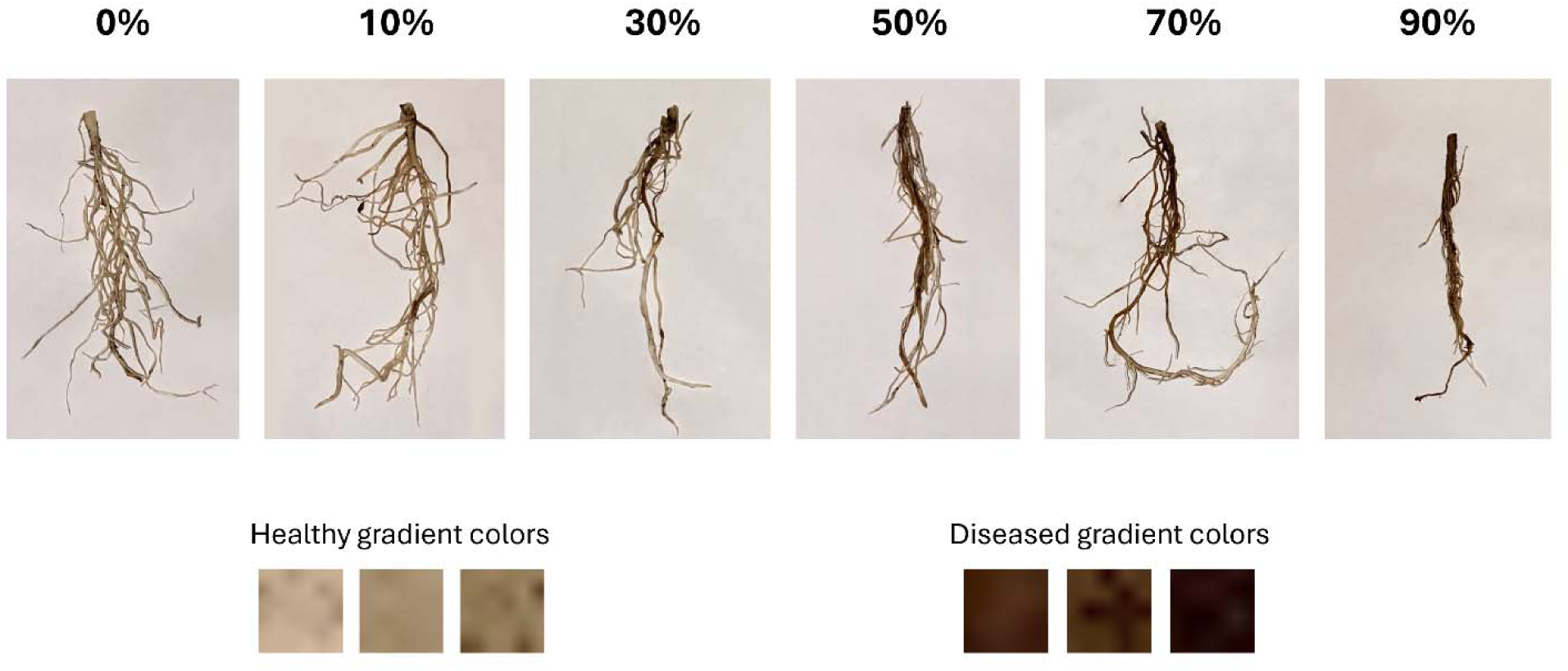
Standard area diagrams (SAD) of soybean roots for visual estimation of root rot severity caused by *Calonectria ilicicola* in a 12-day short-term growth chamber experiment.

#### 3.1.2. SAD validation

The use of SADs significantly improved the accuracy and precision on visual assessment of root rot in controlled conditions (Table 1). Significant increases were observed in Lin’s concordance correlation coefficient (LCCC), the bias correction factor (Cb), and precision (r). Location shift (*u*), and scale shift (*v*) did not differ significantly between aided and unaided assessments.

**Table 1.** Effect of the SAD set on the visual estimation of root rot severity caused by *Calonectria ilicicola*, on the Lin’s concordance correlation coefficient (LCCC, ρc), precision (r), bias coefficient (Cb), location shift parameter (*u*), scale shift parameter (*v*), absolute error, and standard deviation (SD).

| Variable | Mean value |  | Difference between means | (CI 95%)<sup>a</sup> |  |
| --- | --- | --- | --- | --- | --- |
|  | unaided | aided |  |  |  |
| LCCC ($\rho_c$) | 0.739 | 0.842 | 0.103 | (0.044 ; 0.151) | * |
| Precision (r) | 0.825 | 0.867 | 0.042 | (0.014 ; 0.067) | * |
| Cb | 0.887 | 0.971 | 0.084 | (0.035 ; 0.126) | * |
| $v$ | 1.016 | 0.94 | -0.076 | (-0.156 ; 0.023) | |
| $u$ | -0.043 | -0.084 | -0.041 | (-0.233 ; 0.176) | |
| Absolute error | 15.156 | 11.411 | -3.745 | (-5.107 ; -2.262) | * |
| SD | 15.684 | 14.277 | -1.407 | (-2.078 ; -0.650) | * |

<sup>a</sup> 95% confidence intervals were obtained using 2,000 bootstrap resamples of raters. Differences were considered significant when the confidence interval did not include zero (\*)

SAD use also reduced both absolute error and the variability of estimates among raters. Absolute error was consistently lower with SAD-assisted, particularly at intermediate and high root disease severities, indicating improved estimation accuracy across the severity range (Figure 2).

**Figure 2.**
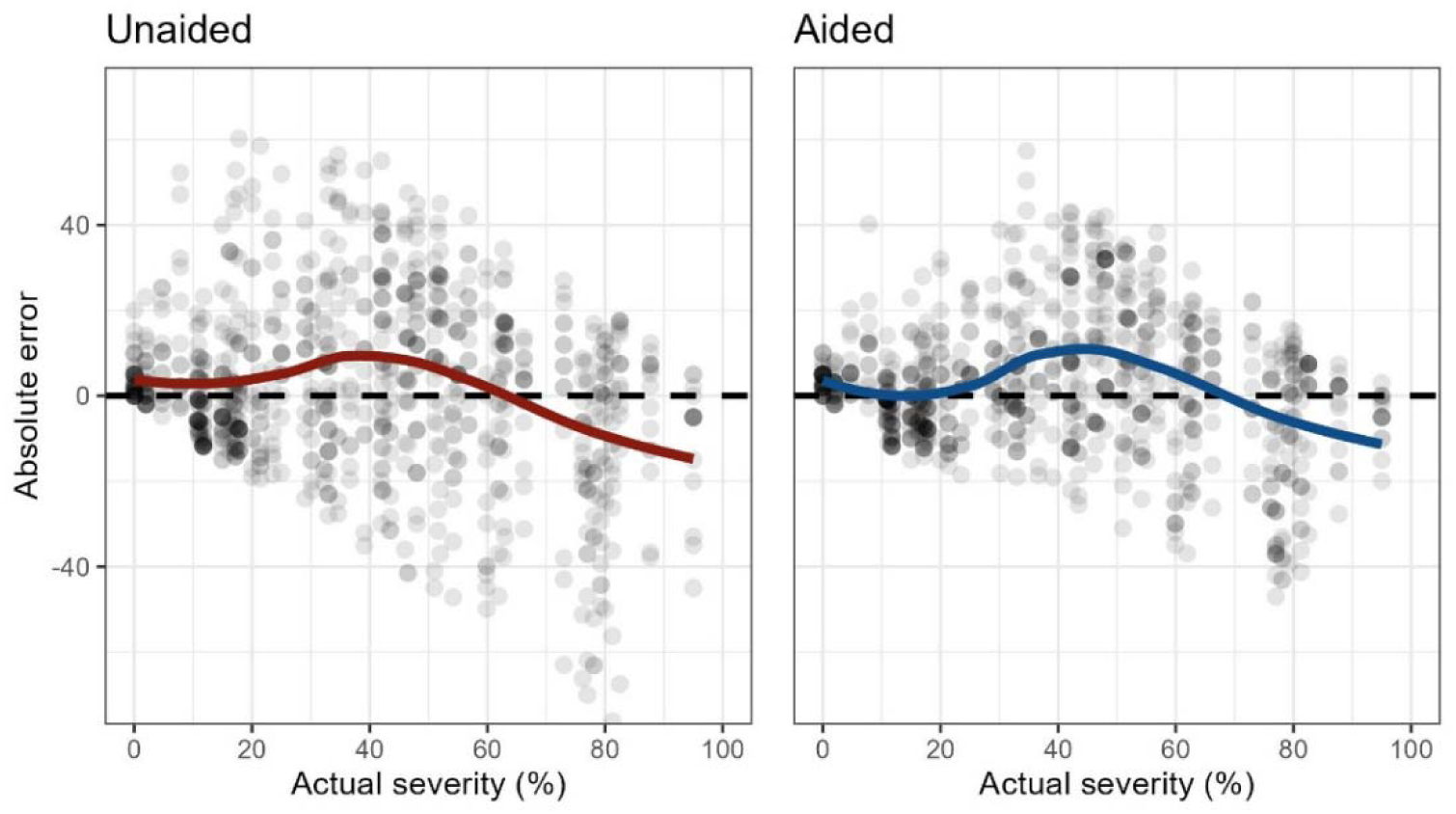
Scatter plot representing the relationship between the raters’ absolute error and actual root necrosis severity (%) under unaided and SAD-assisted assessments. Each point represents an individual observation. Solid lines show the smoothed trend in absolute error across the severity gradient, fitted using the default geom_smooth() function in ggplot2 R package.

Most raters showed gains in accuracy when using the SAD (Figure 3a). Only two raters showed slight reductions in accuracy. However, both had high concordance values even without the SAD, with LCCC decreasing from 0.88 to 0.87 and from 0.91 to 0.90, respectively. In general, the magnitude of the accuracy gain was inversely related to unaided accuracy, with the largest improvement being 0.50 when the unaided LCCC was 0.36. Overall, the use of the SAD significantly increased concordance compared with unaided assessments (Figure 3b).

**Figure 3.**
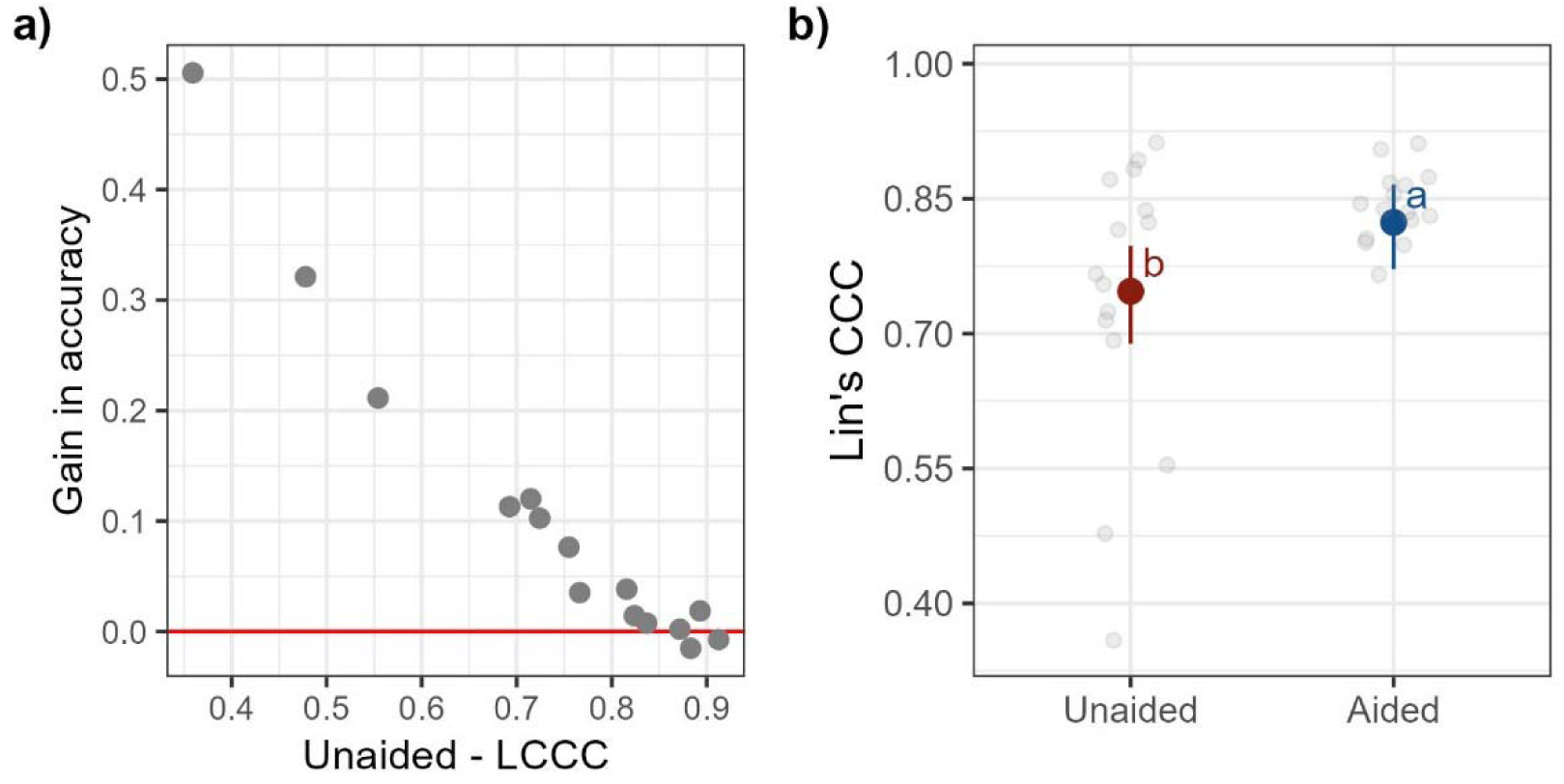
**a)** Relationship between unaided overall accuracy (Lin’s concordance correlation coefficient, LCCC) and the gain in accuracy achieved using SAD set. Each point represents the accuracy gain (difference between SAD-aided and SAD-unaided) for an individual rater. The red line indicates zero gain, separating accuracy improvements (above the line) from accuracy losses (below the line). **b)** Lin’s concordance correlation coefficient (LCCC) of raters’ estimates of root rot severity under unaided and SAD-assisted assessments. Different letters indicate significant differences (Tukey’s HSD test, *p* < 0.05). Bars represent means ± 95% confidence intervals, grey points represent individual raters (n = 15).

#### 3.1.3. Reliability and agreement of root assessments

Among the evaluated methods, the Jiang scale showed the lowest ICC, indicating the lowest inter-rater reliability (Table 2). Use of the SAD increased ICC compared with unaided assessments, indicating greater consistency among raters. OCCC also increased with SAD-assisted assessments, indicating improved overall agreement among evaluators.

**Table 2.** Intraclass correlation coefficient (ICC) and overall concordance correlation coefficient (OCCC) for unaided, SAD-assisted, and Jiang score-based assessments for root rot caused by *C. ilicicola*.

| Assessment method | ICC (95% CI) | OCCC |
| --- | --- | --- |
| Unaided | 0.774 (0.706 ; 0.838) | 0.5135 |
| SAD-assisted | 0.853 (0.803 ; 0.897) | 0.656 |
| Jiang score | 0.74 (0.666 ; 0.812) | - |

To evaluate the ability of each method to discriminate among treatments, three statistically distinct groups were grouped: A, B, and C (Figure 4a). Raters’ performance was classified according to the ability to assign treatments to distinct statistical groups as full discrimination (three groups: all treatments assigned to different groups), partial discrimination (two groups: one treatment was distinguished from the other two), and no discrimination (one group: all treatments belonged to the same group). The SAD-assisted assessment showed the highest discriminatory ability, with 13 raters identifying three distinct groups, compared with unaided assessments, where 6 raters achieved full discrimination and 9 identified two groups. For the ordinal scale, 5 raters achieved full discrimination, 8 identified two groups, and 2 were unable to differentiate among treatments (Figure 4b).

**Figure 4.**
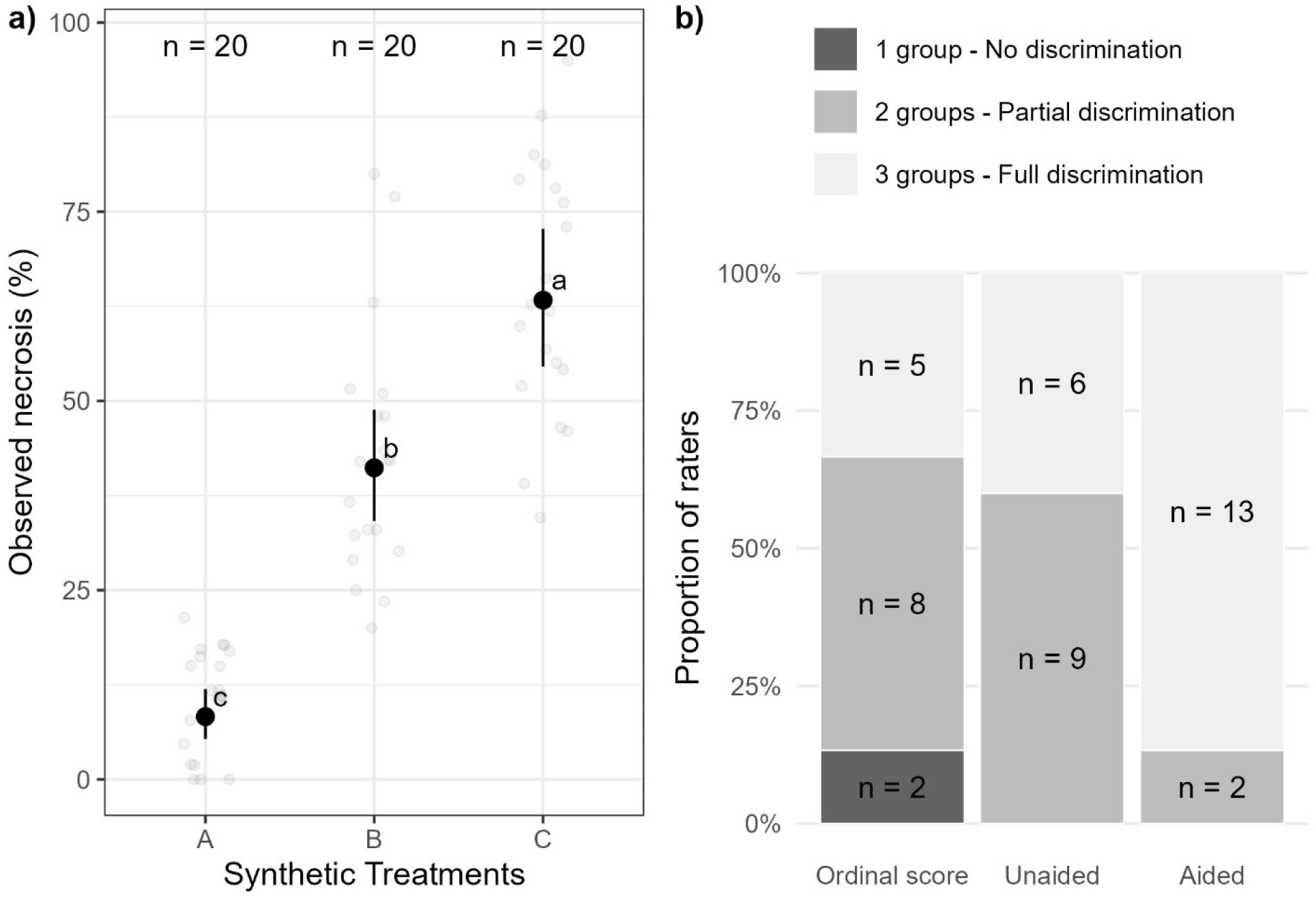
**a)** Distribution of observed necrosis values from images grouped into three synthetic treatments corresponding on three statistical groups (Tukey’s HSD test, *p* < 0.05), bars represent means ± 95% confidence intervals, grey points are observations (n = 20, each treatment). **b)** Proportion and number of raters achieving discrimination among synthetic treatment groups based on ordinal score, unaided and SAD-aided assessment methods. Raters were classified as showing no discrimination (all groups assigned the same category), partial discrimination (two distinguishable groups), or full discrimination (all three synthetic treatment groups correctly distinguished). Ordinal scores were analyzed by proportional odds logistic regression based on ordinal root necrosis rating, and unaided and aided using a generalized linear mixed (Tukey’s HSD, *p* < 0.05).

### 3.2. DSS for canopy disease severity

#### 3.2.1. RCR-DSS development and validation

The RCR-DSS consisted of 10 severity classes ranging from 0 to 9, each defined by specific symptom criteria based on the extent of foliar chlorosis, necrosis, wilting, and plant death (Figure 5a). Representative soybean images were selected for each severity class under natural field conditions to provide a visual reference for symptom expression (Figure 5b).

**Figure 5.**
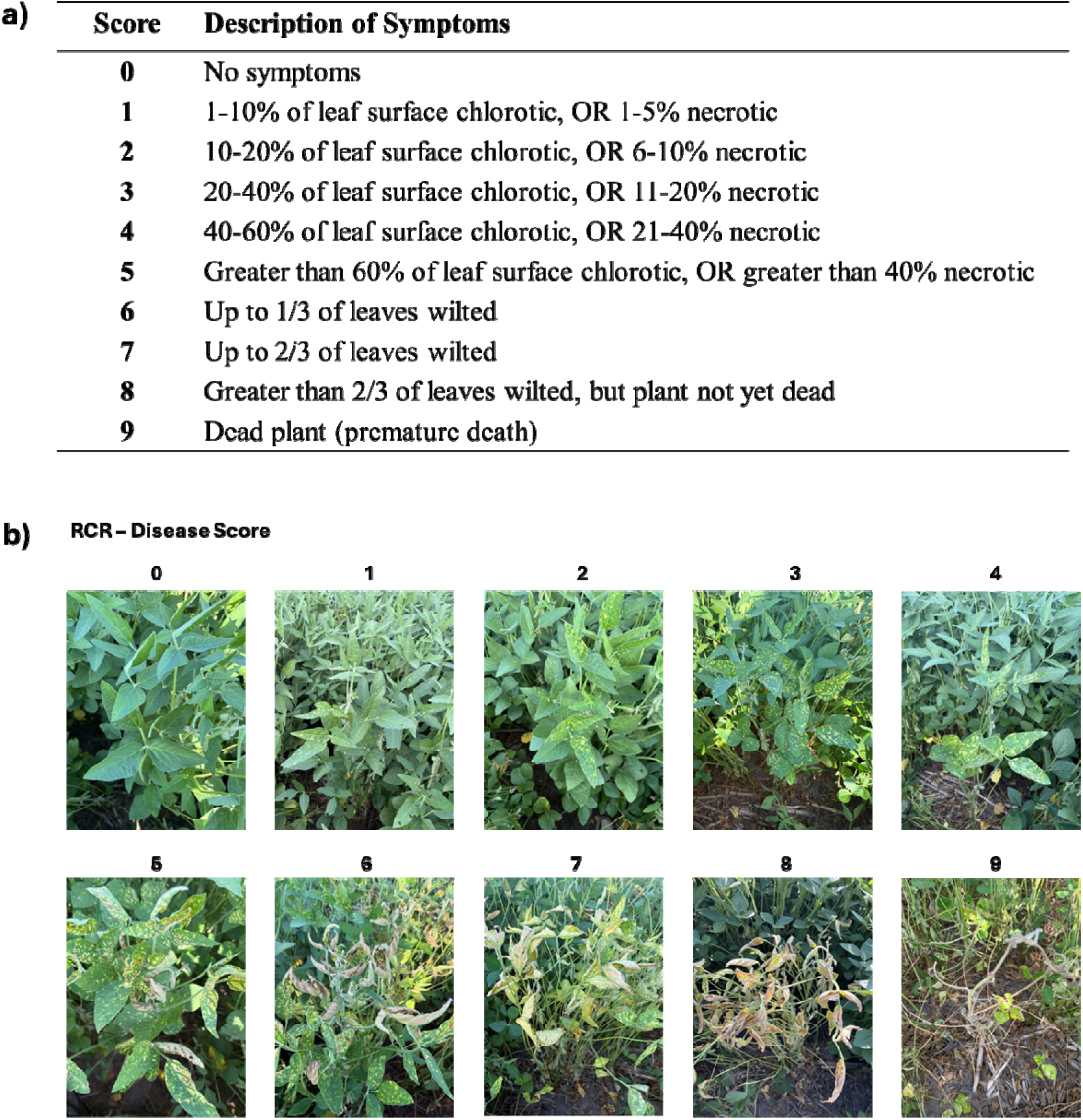
**a)** Symptom descriptions for each red crown rot (RCR) severity score (0–9), caused by *Calonectria ilicicola*. **b)** Diagrammatic symptom scale for RCR (RCR-DSS) severity under field conditions at soybean growth stages R4-R5.

The proposed RCR-DSS showed strong performance across measures of agreement, accuracy, repeatability, and inter-rater reproducibility (Table 3). Lin’s concordance correlation coefficient (LCCC) indicated high agreement between individual rater scores and the consensus reference, with a pooled mean of 0.96 ± 0.03. Mean error values were close to zero across both evaluation rounds, with a pooled value of −0.09 ± 0.32, indicating minimal systematic deviation from the consensus. Individual raters showed both positive and negative deviations from the consensus, suggesting a balanced pattern of slight over- and underestimation. RMSE values indicated generally low overall deviation from the consensus, with a pooled mean of 0.82 ± 0.20.

**Table 3.** Validation metrics of RCR diagrammatic symptom scale (RCR-DSS) performance across evaluation rounds, including intra- and inter-rater reliability.

**Agreement and error-based performance**
| Metrics<sup>a</sup> | Round 1 | Round 2 | Pooled |
| --- | --- | --- | --- |
| Lin's CCC | $0.96 \pm 0.02$ | $0.96 \pm 0.03$ | $0.96 \pm 0.03$ |
| RMSE (accuracy)<sup>b</sup> | $0.77 \pm 0.18$ | $0.84 \pm 0.28$ | $0.82 \pm 0.20$ |
| Mean error (systematic error)<sup>b</sup> | $-0.12 \pm 0.34$ | $-0.06 \pm 0.34$ | $-0.09 \pm 0.32$ |

<sup>a</sup> Values are mean ± standard deviation across raters ( $n = 18$ ). Lin's concordance correlation coefficient (CCC), root mean squared error (RMSE), and mean error were calculated per evaluation round and for the pooled dataset.
<sup>b</sup> No significant differences were inferred, as 95% confidence intervals included zero (2,000 bootstrap resamples)

**Intra-rater repeatability**
| Metric | Min | Max | Mean ± SD |
| --- | --- | --- | --- |
| | 0.888 | 0.995 | $0.96 \pm 0.027$ |

**Inter-rater reproducibility**
|  | ICC (95% CI) |
| --- | --- |
| Round 1 | 0.947 (0.923–0.966) |
| Round 2 | 0.932 (0.903–0.956) |
| Pooled | 0.939 (0.920–0.955) |

Performance varied among individual raters (Figure 6). RMSE values ranged from 0.5 to 1.25 severity units, whereas the mean errors ranged from −0.9 to 0.3. Deviations from the group consensus were generally limited, with most raters remaining within approximately one severity unit of the consensus scoring pattern. Overall, these results indicate close alignment of individual ratings with the group-level reference, despite some variation in rater-specific accuracy and scoring tendency.

**Figure 6.**
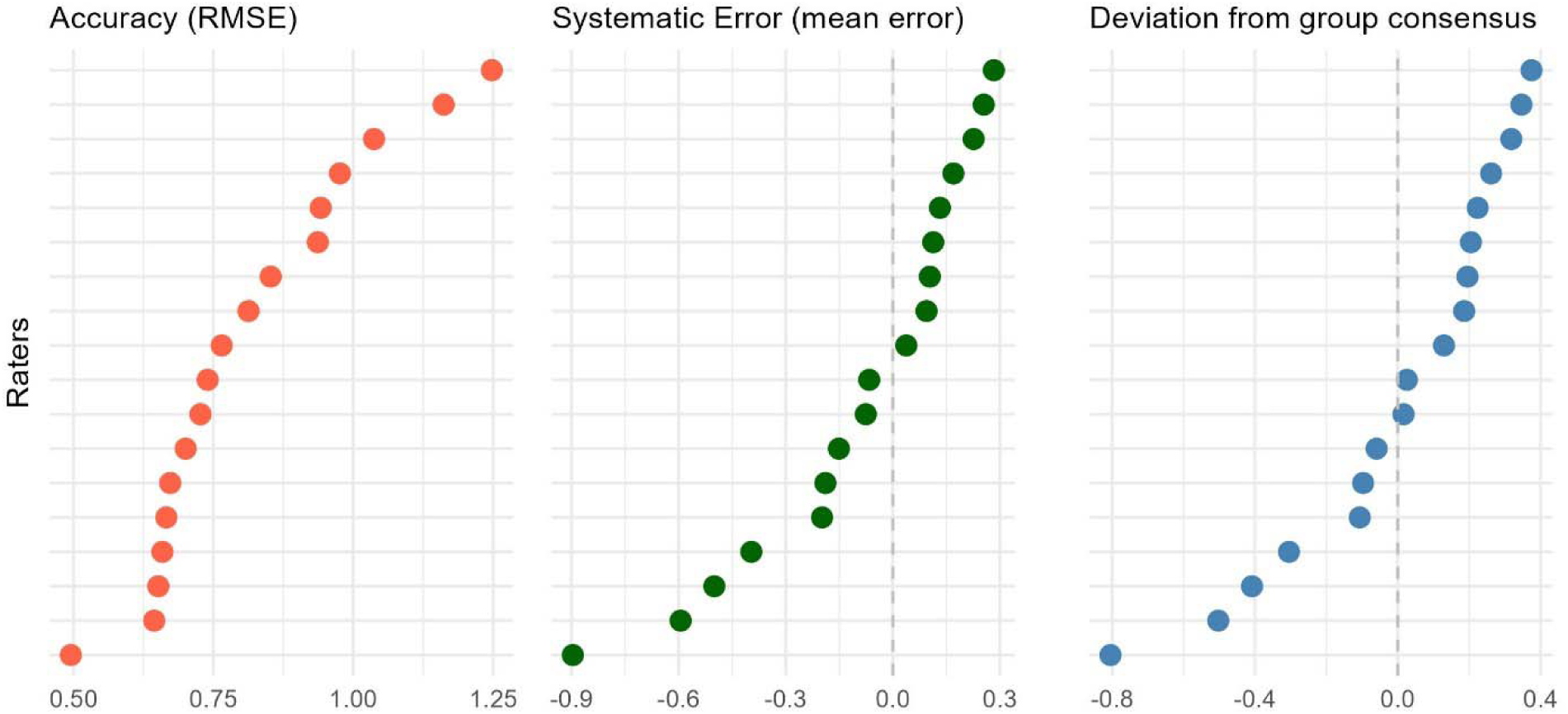
Performance of individual raters in the RCR-DSS validation, expressed as accuracy measured by the root mean square error (RMSE), systematic error (mean error), and deviation from group consensus per picture. Dashed lines indicate references values: no bias in the systematic error, and no deviation from group consensus.

Intra-rater repeatability was high, with quadratic weighted Cohen’s kappa values ranging from 0.888 to 0.995 and a mean of 0.960 ± 0.027, indicating strong consistency within raters across evaluation rounds. In addition, bootstrap confidence intervals for differences in RMSE, absolute error, and LCCC between the two rounds included zero, indicating no evidence of differences in performance between evaluation rounds. Inter-rater reproducibility was also high, with ICC values of 0.947 in Round 1, 0.932 in Round 2, and 0.939 for the pooled dataset, demonstrating excellent agreement among raters.

## 4. Discussion

Quantifying soilborne diseases remains challenging because disease development is expressed across multiple biological scales, from belowground infection to canopy decline, while most assessment methods capture only a single component of this process. RCR is a complex soilborne disease for which both disease severity assessment and crop impact quantification remain challenging due to spatial heterogeneity, temporal dynamics of infection, and overall symptom development. This study addresses these constraints by developing complementary assessment tools from controlled to field conditions, linking root-level quantification and standardized foliar severity assessment in an integrated framework. This multi-scale approach bridges visual disease evaluation with objective measurements, enabling a more consistent and biologically meaningful assessment of RCR development on soybeans.

Although tools based on digital image analysis and artificial intelligence are increasingly available, visual assessment remains the most widely used approach due to its speed, simplicity, and low cost (Bock et al. 2020, 2021; Del Ponte et al. 2017). In this context, diagrammatic scales have become one of the most adopted tools to improve the quality of visual disease assessments. These scales consist of a series of images representing different levels of severity and serve as reference standards during evaluation, helping to reduce observer subjectivity and improve the precision, accuracy, and reproducibility of estimates (Del Ponte et al. 2017, 2021).

*C. ilicicola* colonizes soybean roots and induces necrosis, providing a measurable symptom for quantitative assessment. To our knowledge, no formally validated SADs for quantifying root necrosis percentage are available in existing SAD libraries (Del Ponte et al. 2017; SADBank: http://emdelponte.github.io/sadbank/). This is likely due to difficulties in standardizing root washing procedures, variation in tissue discoloration, the three-dimensional structure of root systems, and challenges in obtaining consistent image-based estimates of diseased tissue (Bock et al. 2010; Cazón et al. 2026). However, under short-term controlled conditions (e.g., ∼12-day-old roots), *C. ilicicola* infection produces a distinct and consistent coloration pattern that allows accurate measure of necrotic tissue (Kleczewski and Geisler 2022). Thus, roots from controlled-condition experiments provide a valuable platform for evaluating product efficacy under uniform and reproducible conditions, enabling consistent comparisons across treatments (Paredes et al. 2026). Given that *C. ilicicola* induces a consistent root necrosis pattern, the SAD presented in this study was developed using true-color imagery to capture these differences more accurately. True-color formats have recently been increasingly adopted in plant disease assessment to more accurately represent disease symptoms as they are observed under field conditions (Del Ponte et al. 2017). This approach aligns with previous evidence from Franceschi et al. (2020), who demonstrated that, for soybean rust, photographic SADs based on true-color images improved rater accuracy and inter-rater reliability compared with grayscale versions.

The effect of specimen size on the accuracy of plant disease estimates has not been fully investigated (Bock et al. 2010). Nita et al. (2003) reported that estimation error increased with leaf size, and Hilton et al. (2024) similarly found higher accuracy and lower bias for small compared with large leaflets, consistent with magnitude-related bias. A comparable source of variation may occur in root systems, where size changes with plant age could influence visual estimates of necrosis. Therefore, our validation was restricted to uniform 12-day-old roots under controlled conditions to standardize root size and minimize this potential effect, as older roots generally develop larger root systems.

Previous research has explored both ordinal and continuous approaches for disease severity assessment (Bock et al. 2021; Chiang and Bock 2021; Del Ponte et al. 2017). Although RCR severity has traditionally been assessed using ordinal root damage scales with four to six categories (Akamatsu et al. 2020; Jiang et al. 2020; Nishi et al. 1999), percent root necrosis has also been used as a continuous measure (Kleczewski & Geisler, 2022). However, no study has directly compared these approaches for treatment discrimination. In our study, we compared root necrosis expressed as continuous percentage estimates with the categorical scoring system proposed by Jiang et al. (2020), finding that percentage-based assessment performed better in discriminating treatment effects. This supports previous evidence that percentage-based scales can reduce estimation error and increase sensitivity compared with ordinal rating systems (Chiang and Bock 2021; Hartung and Piepho 2007). Overall, our results support the use of direct percentage estimation whenever feasible due to improved measurement accuracy and treatment separation.

In continuous assessments, raters often show a preference for rounding severities at intervals of 5% and particularly 10% (Bock et al. 2008; Schwanck and Del Ponte 2014), which may introduce systematic error (Bock et al. 2020). A similar pattern was observed in our study, although raters were allowed to use 1% increments, most scores were recorded in 5% steps. Despite this rounding behavior, performance metrics remained consistent, indicating that the SAD framework is robust under practical scoring constraints. Lin’s concordance correlation coefficient (LCCC), a benchmark metric for comparing rater estimates with actual disease severity (Bock et al. 2021; Del Ponte et al. 2017), showed that the proposed SAD improved overall agreement, accuracy, precision, and bias compared with unaided assessments. To our knowledge, this represents the first validated SAD for quantifying root necrosis percentage in root-affected pathosystems.

Foliar symptoms of RCR typically develop during reproductive stages under field conditions (Hartman et al. 2015). The complex expression may be associated with the distal transport of *C. ilicicola* phytotoxins rather than direct fungal colonization of the leaf tissues (Ochi et al. 2011). Consequently, canopy symptoms reflect the physiological response of the plant rather than direct tissue colonization, making quantitative assessment more challenging. Additionally, RCR-like symptoms may be confused with those caused by other diseases, particularly SDS (Kleczewski et al. 2023). Reliable and standardized evaluation of foliar symptoms is therefore essential for detecting treatment differences, evaluating host resistance, and supporting breeding programs (Jiang and Xie 2023).

Because RCR foliar symptoms are expressed as a complex combination of chlorosis, interveinal necrosis, and wilting or defoliation, they cannot be accurately represented as a continuous percentage of affected leaf area. Therefore, the proposed RCR-DSS was developed as a descriptive ordinal scale, adapted from a 9-point SDS severity scale, to capture the progression of symptom expression (Njiti et al. 1996). Unlike traditional validation approaches that compare unaided and aided assessments, the present validation focused on the reliability, repeatability, and reproducibility of the newly developed RCR-DSS (Madden et al. 2017; Nutter et al. 1993). Consequently, all raters used the scale in both evaluation rounds. Although the absence of unaided assessments precluded direct quantification of the improvement provided by the scale (Del Ponte et al., 2017), the primary objective was to determine whether the RCR-DSS generated consistent severity estimates within and among evaluators. The proposed RCR-DSS demonstrated high reliability, repeatability, and accuracy, supporting its suitability as a standardized tool for assessing RCR severity under field conditions.

## Conclusion

The integration of root-level necrosis quantification and standardized foliar severity assessments provides a robust framework for assessing RCR development across experiments. Each method has inherent strengths and limitations: root assessments provide direct measures of belowground damage but are destructive and labor-intensive; visual foliar ratings capture disease-specific symptom expression but are subjective and limited in spatial coverage. By combining these complementary sources of information, the multiscale approach improves the consistency and biological relevance of disease assessment. Overall, this integrated framework enhances the ability to characterize RCR dynamics and supports a more comprehensive evaluation of disease impacts.

